# Neutrophil and monocyte dysfunctional effector response towards bacterial challenge in critically-ill COVID-19 patients

**DOI:** 10.1101/2020.12.01.406306

**Authors:** Srikanth Mairpady Shambat, Alejandro Gómez-Mejia, Tiziano A. Schweizer, Markus Huemer, Chun-Chi Chang, Claudio Acevedo, Judith Bergada Pijuan, Clement Vulin, Nataliya Miroshnikova, Daniel A. Hofmänner, Pedro D. Wendel Garcia, Matthias P Hilty, Philipp Bühler Karl, Reto A. Schüpbach, Silvio D. Brugger, Annelies S. Zinkernagel

**Affiliations:** Department of Infectious Diseases and Hospital Epidemiology, University Hospital of Zurich, University of Zurich, Switzerland; Institute of Intensive Care, University Hospital of Zurich, University of Zurich, Switzerland

**Keywords:** COVID-19, Neutrophils, Monocytes, secondary bacterial infections, hypercytokinemia

## Abstract

COVID-19 displays diverse disease severities and symptoms. Elevated inflammation mediated by hypercytokinemia induces a detrimental dysregulation of immune cells. However, there is limited understanding of how SARS-CoV-2 pathogenesis impedes innate immune signaling and function against secondary bacterial infections. We assessed the influence of COVID-19 hypercytokinemia on the functional responses of neutrophils and monocytes upon bacterial challenges from acute and corresponding recovery COVID-19 ICU patients. We show that severe hypercytokinemia in COVID-19 patients correlated with bacterial superinfections. Neutrophils and monocytes from acute COVID-19 patients showed severely impaired microbicidal capacity, reflected by abrogated ROS and MPO production as well as reduced NETs upon bacterial challenges. We observed a distinct pattern of cell surface receptor expression on both neutrophils and monocytes leading to a suppressive autocrine and paracrine signaling during bacterial challenges. Our data provide insights into the innate immune status of COVID-19 patients mediated by their hypercytokinemia and its transient effect on immune dysregulation upon subsequent bacterial infections

## Introduction

While most patients with Coronavirus-disease-2019 (COVID-19) exhibit only mild to moderate symptoms, approximately 10% to 15% of patients progress to a severe disease. This severe course of COVID-19 may require intensive care unit (ICU) support (Wu and McGoogan, 2020) and is characterized by acute respiratory distress syndrome (ARDS) as well as cardiovascular, gastrointestinal and neurological dysfunctions (Guan et al., 2020; Shi et al., 2020; The, 2012; Wu et al., 2020; Zhou et al., 2020).

Despite limited data, bacterial superinfections in COVID-19 pneumonia (Hughes et al., 2020; Lansbury et al., 2020), contribute to mortality (Chen et al., 2020; He et al., 2020; Zhou et al., 2020). In our recent prospective single centre cohort study, we showed that 42.2% of the ICU COVID-ARDS patients had bacterial superinfections (Buehler et al., 2020). These were associated with reduced ventilator-free survival and significantly increased ICU length of stay (LOS) (Buehler et al., 2020).

Beyond ARDS, COVID-19 patients have been reported to show a complex immune dysregulation, characterized by misdirected host responses and altered levels of inflammatory mediators (Chen et al., 2020; Giamarellos-Bourboulis et al., 2020b; Wang et al., 2020; Wen et al., 2020). Severe COVID-19 is characterized by lymphopenia, neutrophilia and myeloid cell-dysregulation (Kuri-Cervantes et al., 2020; Tan et al., 2020; Wang et al., 2020; Wen et al., 2020; Wu et al., 2020) as well as high plasma cytokine levels (Arunachalam et al., 2020; Lucas et al., 2020). These high cytokine levels have been suggested to result in functional paralysis of the immune cells, causing respiratory and multiple organ failure (Giamarellos-Bourboulis et al., 2020b). However, there is a limited understanding of how SARS-CoV-2 pathogenesis impedes innate immune signaling and function against secondary bacterial infections. Similarly, the role of neutrophils and monocytes and their ability to respond to bacterial infection during COVID-19 remains to be elucidated. Here, we sought to investigate the functional response of neutrophils and monocytes derived from critically-ill COVID-19 patients during their acute illness and their subsequent recovery (rec)-phase towards bacterial challenge as well as the signaling mediators underlying this response.

## Results

### Extensive COVID-19-mediated hypercytokinemia correlates with subsequent bacterial superinfections

We first assessed the plasma levels of cytokines involved in neutrophil and monocyte functional responses in our prospective cohort of critically-ill COVID-19 ICU patients (acute, n=27), including the same patients in their recovery phase (rec, n=21), as well as healthy donors (n=16) (Table S1 and S2). As shown previously (Blanco-Melo et al., 2020; Huang et al., 2020; Lucas et al., 2020) we observed that the cytokines affecting neutrophil function granulocyte colony-stimulating factor (G-CSF), interleukin (IL)-8, IL-4, macrophage inflammatory protein (MIP-1α, MIP-2α, MIP-1β) and stromal cell-derived factor 1 alpha (SDF-1α) had significantly increased levels in acute-patients. In addition, we showed that these levels decreased upon recovery and were similar to values measured in healthy donors (Fig. S1). For monocyte effectors, we found the most significant changes in the levels of fractalkine (CX_3_CL1), interferon gamma-induced protein 10 (IP10) and monocyte chemotactic protein-1 (MCP-1) (Fig. S1).

In a next step, we sought to investigate whether the cytokine levels varied in COVID-19 patients who developed secondary bacterial infections as compared to patients who did not. Principal component analysis (PCA) showed that cytokines measured in COVID-19 patients clustered apart from healthy donors (Fig. 1A). Both, acute- (Fig. 1A, right top panel) and rec-phase COVID-19 patients (Fig. 1A, right bottom panel), who developed a secondary bacterial infection displayed higher degrees of hypercytokinemia with increased separation on the density curve as compared to those without (Fig. 1A). This was confirmed by calculating the normalised cytokine values (sum of Z-scores) in the plasma. Patients who developed a secondary bacterial infection showed significantly elevated cumulative cytokine levels in both acute- and rec-phase (Fig. 1B). Additionally, we found a distinct clustering of specific cytokines among critically-ill COVID-19 patients who developed bacterial superinfections versus patients without any bacterial superinfections (Fig. S2A-B). Furthermore, integrative correlation mapping of clinical parameters taken within 24 hours from sampling revealed that cytokine levels correlated with myoglobin levels and bacterial superinfection status, which in turn, correlated with ICU LOS and ventilation days (Fig. 1C) (Buehler et al., 2020). Overall, extensive COVID-19 hypercytokinemia correlated with the development of bacterial superinfections.

**Figure 1.**
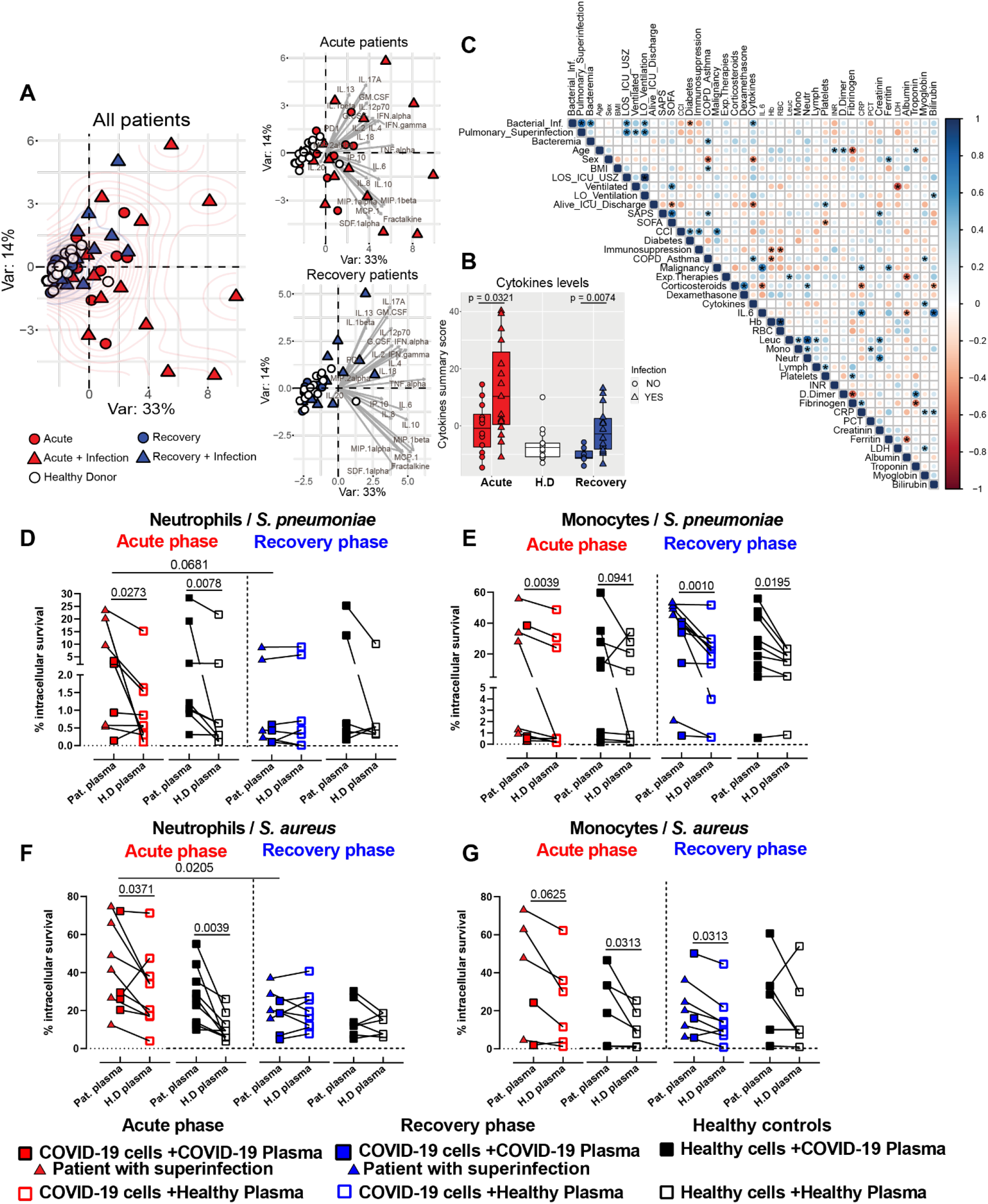
Characterization of inflammatory mediators in COVID-19 plasma and impaired bactericidal capacity of innate immune cells. (**A**) PCA of healthy donors (white) vs acute (red) and rec (blue)-COVID-19 patients grouping the plasma cytokine levels and status of secondary bacterial infections. Patients with secondary bacterial infection are depicted as triangle and patients without superinfection as circle symbols. (**B**) Normalised cytokine values (sum of Z-scores) in the plasma of acute (red), rec (blue) patients with or without bacterial superinfection and healthy donors (white) (**C**) Integrated correlation clustering map of relevant clinical parameters; circle-color indicates positive (blue) and negative (red) correlations, color intensity represents correlation strength as measured by the Pearson’s correlation coefficient. **(D-E)** Intracellular killing capacity of COVID-19 patient (acute-red and rec-blue) neutrophils (left) and monocytes (right) (n=8-10) pre-exposed to plasma from patients (solid symbols) vs healthy plasma (open symbols) upon infection with SP (**D**) or SA (**E**).

### Reduced elimination of intracellular bacteria by neutrophils and monocytes in acute-phase COVID-19 patients

These above described clinical findings indicated increased susceptibility towards bacterial superinfection in critically-ill COVID-19 patients associated with alterations in plasma cytokine levels. Aiming to further dissect these findings, we assessed neutrophil and monocyte function upon bacterial challenge *ex vivo*. Neutrophils and monocytes derived from critically-ill COVID-19 patients or healthy donors were incubated with either autologous or heterologous plasma prior to bacterial challenge with *Streptococcus pneumoniae* (SP) or *Staphylococcus aureus* (SA) (Fig. 1D-1G and S2C-F). Neutrophils from acute patients internalized significantly less SP. Stimulation with healthy donor plasma partially restored the internalization ability (Fig. S2C). We did not observe significant differences in the phagocytosis ability of monocytes challenged with SP (Fig. S2D). No plasma-mediated effect on phagocytosis ability was observed when either neutrophils or monocytes were challenged with SA (Fig. S2E-F).

Our data show that acute COVID-19 neutrophils and monocytes had impaired bactericidal function with a significant reduction in their ability to clear bacteria as compared to the same cells stimulated with healthy plasma (Fig. 1D-G). Similarly, stimulation of healthy neutrophils and monocytes with acute plasma showed significantly impaired clearance of intracellular bacteria (Fig. 1D-G). Neutrophils from rec-phase patients did not show any impairment in their ability to eliminate intracellular bacteria compared to healthy cells (Fig. 1D and F). In contrast, monocytes from rec-phase patients still displayed reduced bacterial killing capacity (Fig. 1E and G). Neutrophils and monocytes derived from COVID-19 patients who developed subsequent bacterial superinfections showed a tendency towards decreased intracellular killing capacity compared to COVID-19 patients without (Fig. 1D-G). Further confirmation was achieved by stimulating healthy monocytes with acute plasma, which showed significantly impaired ability to clear intracellular bacteria as compared to monocytes stimulated with rec-phase-patients’ or healthy plasma (Fig. S2G-H). These data suggested that hypercytokinemia during COVID-19 impairs neutrophils’ and monocytes’ ability to eradicate intracellular bacteria.

### Impaired neutrophil and monocyte effector response against bacterial challenges in acute-phase COVID-19 patients

To assess the factors involved in the reduced intracellular killing capacity of acute phase neutrophils, we analyzed key neutrophil effector responses. Neutrophils from acute phase patients stimulated with autologous plasma produced significantly lower levels of reactive oxygen species (ROS) upon bacterial challenge compared to stimulation with healthy plasma (Fig. 2A SA and SP). The same effect was observed when healthy neutrophils were stimulated with acute patients’ plasma prior to bacterial challenge (Fig. 2A). Conversely, neutrophils from rec-patients stimulated with autologous plasma displayed the same ROS production levels as neutrophils derived from healthy donors (Fig. 2B SA and SP).

**Figure 2.**
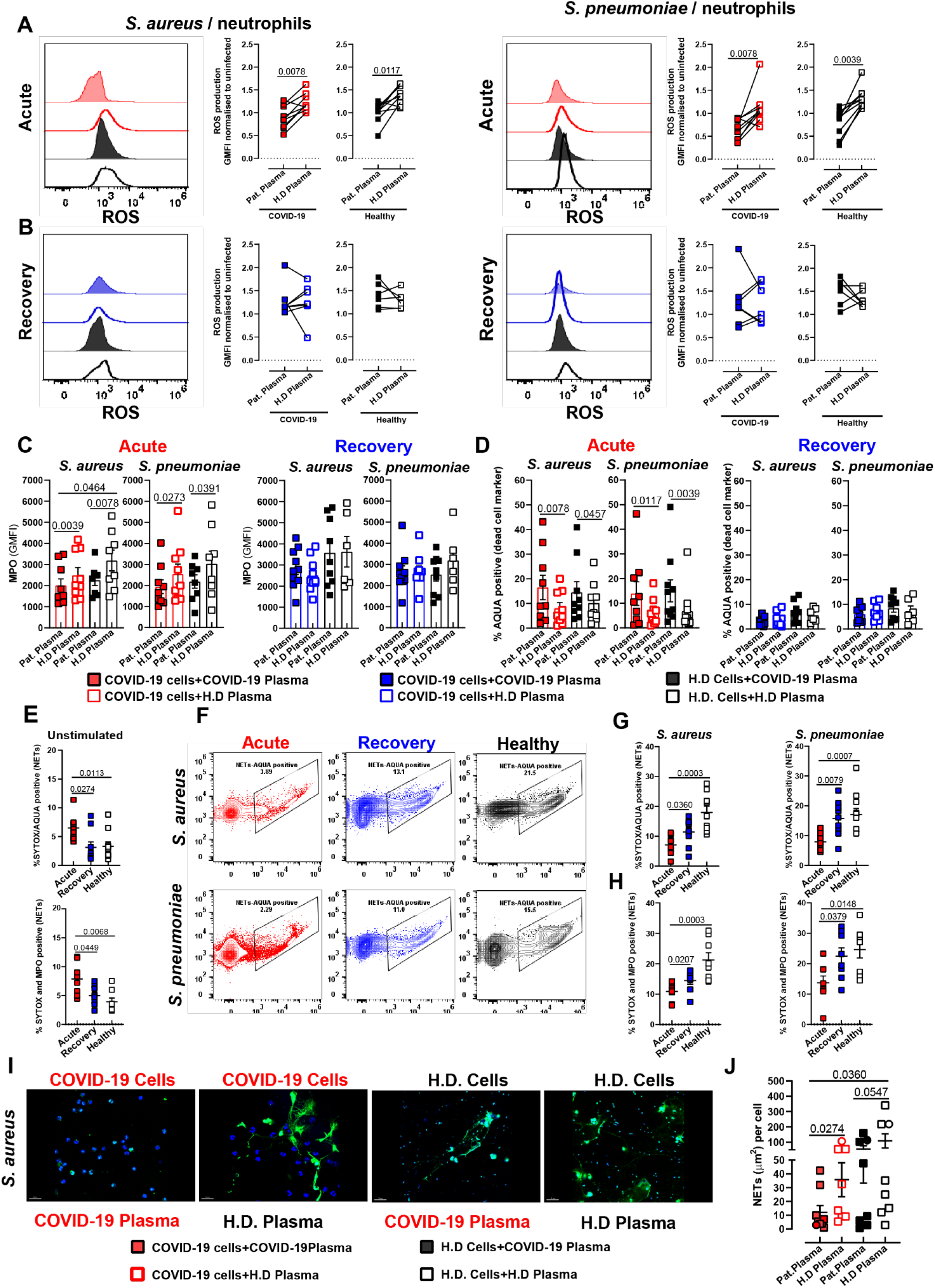
Impaired neutrophil effector response against bacterial challenge in acute COVID-19 patients. Functional characterization of neutrophils pre-exposed to plasma from COVID-19 acute (red) or rec (blue) patients (solid symbols) vs healthy plasma (open symbols) upon challenge with either SA or SP. Neutrophil functionality was assessed by quantification of ROS (**A-acute**) (**B-rec**) (n=7-8), Intracellular MPO (**C**) (right-acute) (left-rec) and cell viability (**D**) (right - acute) (left - rec) (n=7-9). The ability to produce NETs was also measured by flow-cytometry. **E**) SYTOX and AQUA positive cells (top) and MPO-SYTOX positive cells (bottom) under unstimulated COVID-19 conditions. (**F, G and H**) SYTOX and AQUA positive cells (top) and MPO-SYTOX positive cells (bottom) upon bacterial challenge (n=6-8). **I**) Representative confocal images on NETs formation upon challenge with SA using HOECHST (nuclei; blue), SYTOX (staining extracellular-DNA; green). **J**) Quantification of NETs using fluorescence microscopy (squares, n= [226 - 8898]) and confocal microscopy (circles, n= [19 - 113]) nuclei per point (n=7-8).

Additionally, stimulation of neutrophils from acute patients and healthy donors with acute patients’ plasma resulted in significantly lower levels of myeloperoxidase (MPO) compared to stimulation with healthy plasma upon bacterial challenge (Fig. 2C, left). In line with the normalized ROS levels during recovery, neutrophils from rec-phase stimulated with plasma from rec-phase patients exhibited MPO levels comparable to cells stimulated with healthy plasma, after bacterial challenge (Fig. 2C, right). Since increased rates of dysregulated cell death of various cell types during COVID-19 has been described in literature (Varga et al., 2020; Xu et al., 2020), we investigated whether neutrophils from critically-ill COVID-19 patients showed increased sensitivity towards cell death during bacterial infection. Neutrophils stimulated with acute-COVID-19 plasma, irrespective of their origin, showed increased cell death upon bacterial challenge (Fig. 2D, left). In contrast, neutrophils stimulated with plasma from rec patients or healthy donors prior to bacterial challenge remained viable (Fig. 2D, right).

Recently, it has been proposed that neutrophil extracellular traps (NETs) contribute to the formation of microthrombi in COVID-19 ARDS and that sera from COVID-19 patients triggered NETs release in healthy neutrophils (Middleton et al., 2020; Zuo et al., 2020). Since NETs formation is a strategy to eliminate extracellular pathogens (Brinkmann et al., 2004), we tested the hypothesis that bacterial challenge-mediated cell death of neutrophils isolated from acute patients is due to increased NETs release. Neutrophils from acute patients exhibited a higher amount of spontaneous extracellular DNA-release (Fig. 2E, top) and elevated levels of MPO-DNA (Fig. 2E, bottom) than neutrophils from recovery patients or healthy donors. However, bacterial challenge resulted in significantly lower release of NETs and MPO-DNA from neutrophils derived from acute patients as compared to neutrophils from rec-phase or healthy donors (Fig. 2F-H). The inability to release NETs upon bacterial challenge was confirmed by fluorescence microscopy (Fig. 2I and J, Fig. S3B).

Analysis of monocyte subsets revealed significantly lower proportions of classical (CD14+ CD16-) monocytes during acute COVID-19. Similarly, non-classical (CD14dim CD16+) monocytes proportions were reduced during both acute and rec-phase COVID-19 compared to healthy donors (Fig. S4A). We observed the same plasma-mediated decrease of ROS levels upon bacterial challenge in the acute-phase in classical monocytes, whereas no differences in nitric oxide production were found (Fig. S4B-F). Non-classical monocytes exhibited no difference in ROS production, irrespective of disease status (Fig. S4G-H). Together, these data suggest that neutrophils from acute-COVID-19 patients are in a state of exhaustion causing inability to produce ROS, MPO and to trigger NETs release upon secondary bacterial challenge, whereas classical monocytes were skewed towards a significantly impaired ROS, but not nitric oxide, production.

### Neutrophils cell surface receptor alterations in acute COVID-19 patients contribute to an dysfunctional phenotype

Given the observed impaired neutrophil effector response to bacterial challenges in acute COVID-19 patients, we investigated potentially pivotal signaling mechanisms and receptor phenotypes of neutrophils. Neutrophils from acute patients showed a significant decrease in the expression of the receptors CXCR 1, 2, 3, CCR1 and CCR5 (Fig. 3A, B, E, and F) and of the maturation marker CD15 compared to neutrophils during recovery or from healthy donors (Fig. 3D). Additionally, we observed higher levels of the activation marker CD66b and chemokine receptor CXCR4 in neutrophils from acute patients, indicating the presence of immature or dysfunctional neutrophils in the blood. (Fig. 3C and S5D). Furthermore, upon bacterial challenge a similar pattern of cell surface receptor phenotype with reduced expression of CXCR1, 2, 3, CCR1, 5 and CD15, with increased expression of CXCR4 and CD66b was found in neutrophils from acute COVID-19 (Fig. 3A-F and S5C-F).

**Figure 3.**
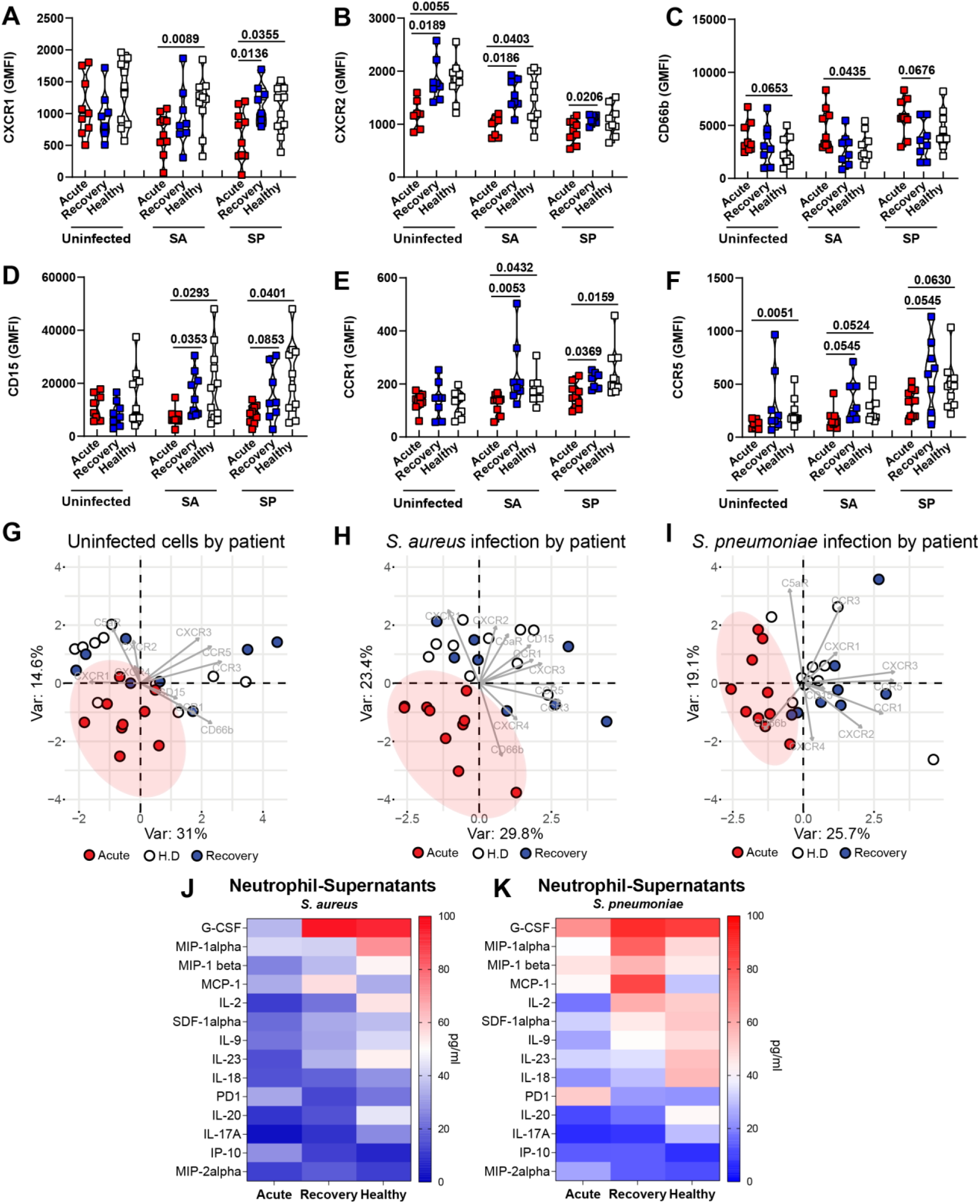
Expression of surface markers and secretion of cytokines in neutrophils upon bacterial challenge. Expression of key surface markers in COVID-19 acute (red), rec (blue) and healthy donors’ (white) neutrophils (**A-F**) (n=8-10). PCA of cell surface phenotype of COVID-19 patients acute (red), rec (blue) and healthy donors’ (white) neutrophils at the basal level without bacterial challenge (COVID-19 status) (**G**), upon SA infection (**H**) or SP infection (**I**). Heat map of cytokine secreted by neutrophils from COVID-19 patients (acute and rec) and healthy controls after bacterial challenge with SA (left) or SP (right) (n=3-4) (**J-K**).

Overall, PCA analysis of acute-COVID-19 patients showed clear separation from rec-phase patients and healthy donors, exhibiting a strong clustering for their receptor phenotype, whereas rec-phase and healthy controls largely overlapped (Fig. 3G-I). Finally, we studied whether this distinct neutrophil phenotype in COVID-19 patients contributed to an impaired cytokine production involved in the autocrine-paracrine signaling upon bacterial challenge. We found that acute COVID-19 neutrophils were characterized by reduced secretion of G-CSF, MIP-1α, MIP-1β, MCP-1, IL-2, SDF-1α, IL-9, IL-17A, IL-18, IL-20 and IL-23, but increased secretion of soluble PD1, IP-10 and MIP-2α compared to rec-phase and healthy neutrophils (Fig. 3J-K). Collectively, these data suggest that acute COVID-19 is marked by the presence of dysfunctional neutrophils, displaying reduced effector responses upon secondary bacterial challenge. Together with plasma cytokine levels affecting neutrophil function (Fig. S1) and the clinical observation that the patients in our prospective cohort presented with neutrophilia, our data helps to explain the role of neutrophil dysfunction in increased risk of secondary bacterial infections in critically-ill COVID-19 patients.

### Monocyte subpopulation alterations in COVID-19 contribute to impaired response against bacterial challenges

The myeloid compartment, especially monocytes, is particularly affected by COVID-19 (Schulte-Schrepping et al., 2020). Classical monocytes from acute patients displayed more heterogeneity, with higher expression of CD163, CX_3_CR1 and low expression of HLA-DR compared to recovery and healthy monocytes (Fig. 4A and C, S6A). Upon bacterial challenge, classical monocytes from COVID-19 patients (acute and recovery) displayed high expression of CD163 and CD11b, but low expression of the activation markers HLA-DR, CD86 and CD80 (Fig. 4A-D, and S6). Overall, PCA analysis showed that the acute COVID-19 classical monocyte clustering pattern was strongly associated with low expression of HLA-DR, CD86, CD80 and a high expression of CD163, CX3CR1 and CD11b (Fig. 4G-I).

**Figure 4.**
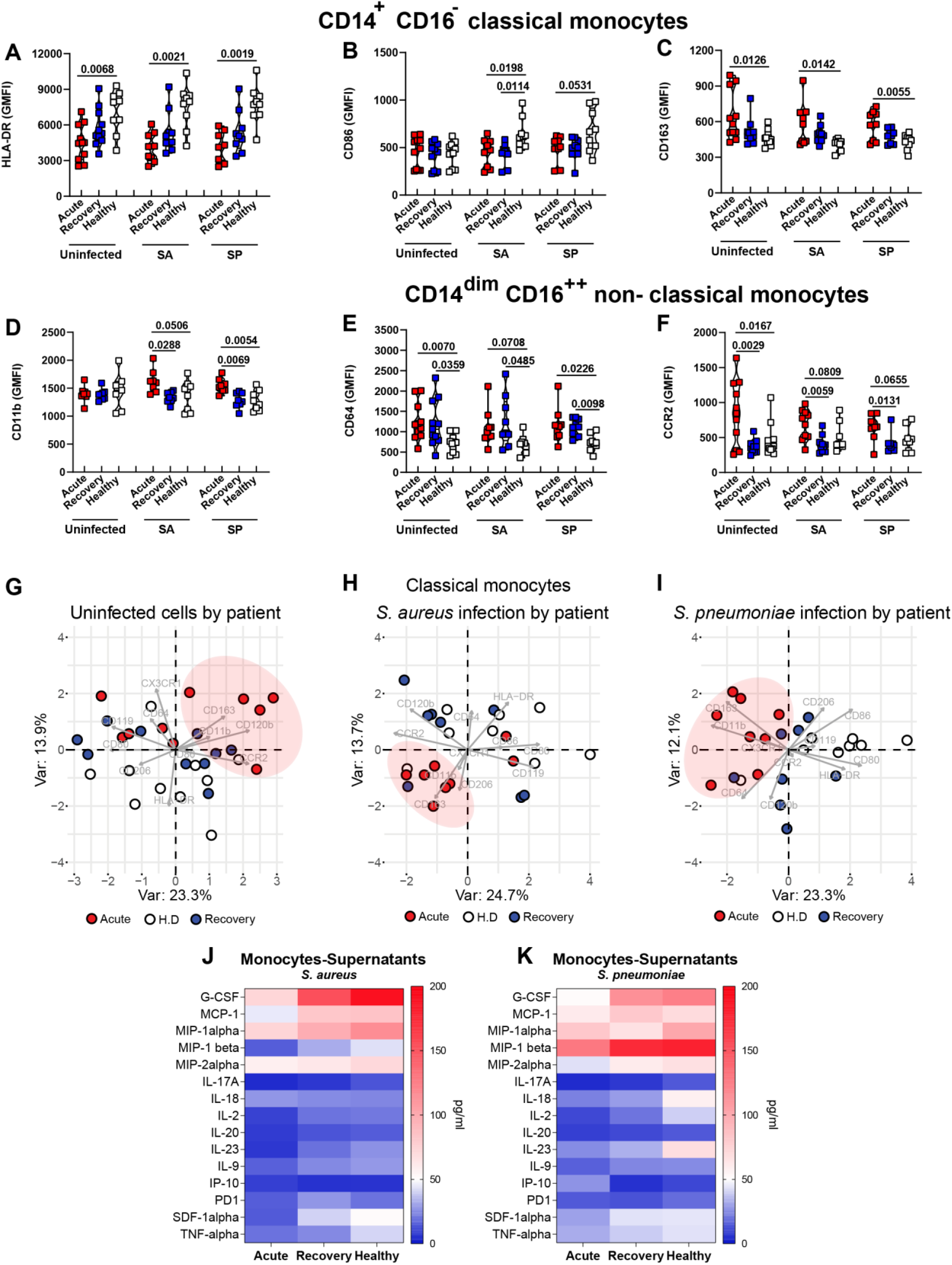
Phenotypic characterization of surface markers and the secretion of cytokines in monocytes upon bacterial challenge. Expression of key surface markers in COVID-19 acute (red), rec (blue) and in healthy donors’ (white) classical (**A-C**) and non-classical (**D-F**) monocytes (n=9-11). PCA of cell surface phenotype of COVID-19 patients acute (red), rec (blue) and in healthy donors (white) of classical monocytes at the basal level without bacterial challenge (COVID-19 status) (**G**), upon SA infection (**H**) or SP infection (**I**). Heat map of cytokine secreted by monocytes from COVID-19 patients (acute and rec) and healthy controls after bacterial challenge with SA (left) or SP (right) (n=2-3) (**J-K**).

Moreover, non-classical monocytes showed higher expression of CCR2, CD11b, CD163 and CD86 in acute-phase, while CD64 was increased in both acute- and rec-phase compared to healthy controls, both in the presence and absence of bacterial challenge (Fig. 4E-F, and S6B). Similarly, PCA analysis showed that the non-classical monocyte cluster in the acute-phase was characterized by an increased expression of CCR2, CD163 CD120b, CD11b and low expression of HLA-DR (Fig. S6C). Finally, COVID-19 monocytes showed a dampened cytokine response to challenge with bacteria compared to healthy controls (Fig. 4J and K). Particularly, monocytes from patients with acute COVID-19 showed reduced secretion of G-CSF, MIP-1α, MIP-1β, MCP-1, TNF-α and IL2 (Fig. 4J and K).

Taken together, these data suggest that dynamic changes of monocyte receptor and cytokine secretion profile associated with acute-COVID-19-were involved in an aberrant antibacterial response.

## Discussion

We show that a higher degree of COVID-19 mediated hypercytokinemia in the plasma is positively associated with bacterial superinfections in COVID-19 patients. Neutrophils and monocytes from acute-phase COVID-19 patients exhibited impaired microbicidal capacity, reflected by abrogated ROS and MPO production as well as NETs formation by neutrophils and impaired ROS production in monocytes. This immunosuppressive phenotype was characterized by a high expression of CD15, CXCR4 and low expression of CXCR1, CXCR2 and CD15 in neutrophils and low expression of HLA-DR, CD86 and high expression of CD163 and CD11b in monocytes. Additionally, neutrophils and monocytes from acute COVID-19 exhibited a blunted cytokine production capacity upon bacterial challenge.

Studies have shown that severe COVID-19 is accompanied by hypercytokinemia with high levels of pro-inflammatory cytokines such as IL-6 and IL-1β as well as anti-inflammatory cytokines such as IL-4 and IL-10 (Arunachalam et al., 2020; Coperchini et al., 2020; Lucas et al., 2020). Our initial screening detected significantly higher levels of cytokines involved in recruitment and trafficking of neutrophils such as IL-8, G-CSF and SDF-1α, in accordance with previous reports showing that acute COVID-19 patients have elevated neutrophil counts (Chevrier et al., 2020; Morrissey et al., 2020; Schulte-Schrepping et al., 2020; Silvin et al., 2020; Wu et al., 2020). We also report a rise in CX_3_CL1, IP10 and MIP-1β levels, indicating increased recruitment of monocytes (Chevrier et al., 2020; Lucas et al., 2020). However, the concomitant presence of high levels of IL-4 and IL-10, with broad anti-inflammatory functions, might cause functional impairment of neutrophils and monocytes towards bacterial challenge (Woytschak et al., 2016). We found that higher degree of hypercytokinemia in the plasma correlated with the occurrence of bacterial superinfections in COVID-19 patients (Buehler et al., 2020). Specifically, TNF-α, IFN-γ, G-CSF, MIP-1α, IL-10 and CX_3_CL1 were elevated in patients developing bacterial superinfection. However, further studies using larger cohorts specifically looking at the correlation between elevated levels of certain cytokines and the risk for developing bacterial superinfections are required to elaborate on these observations.

We hypothesized that elevated levels of inflammatory mediators in the plasma might lead to impaired functional responses to bacterial challenge. Indeed, neutrophils and monocytes derived from acute COVID-19 showed a decreased capacity to kill intracellular bacteria. The capacity to clear internalized bacteria could be restored when COVID-19 derived cells were stimulated with healthy plasma. Additionally, monocytes but not neutrophils, from rec-patients also showed impaired ability to clear intracellular bacteria. We were able to link this inefficiency in clearing intracellular bacteria in acute COVID-19 by a significant decrease in their ability to produce ROS and intracellular MPO (in neutrophils).

This altered functionality is consistent with a recent study showing reduced oxidative burst in response to *E. coli* in severe COVID-19 patients (Schulte-Schrepping et al., 2020). Since neutrophils and monocytes engage in a complex crosstalk with other immune cells to elicit efficient effector response, we were keen on identifying possible autocrine-paracrine signaling mechanisms. Neutrophils and monocytes from critically-ill COVID-19 patients were functionally impaired in their capacity to produce cytokines important for activation and subsequent antimicrobial actions. Significantly lower levels of G-CSF and IL-17 as well as IL-18 in neutrophils from acute patients could be linked to decreased ROS (Castellani et al., 2019; Hu et al., 2017) and MPO (Leung et al., 2001) production, respectively. This was consistent with a recent observation regarding the diminished or inexistent expression of cytokine genes (IL6, TNF-α) by monocytes upon stimulations with TLR ligands (Arunachalam et al., 2020). Several recent studies have proposed that NETs can contribute to inflammation-associated lung damage and microthrombi in severe COVID-19 patients (Middleton et al., 2020; Radermecker et al., 2020). The concentration of NETs components was found to be augmented in plasma, tracheal aspirate and lung autopsy tissues from COVID-19 patients (Veras et al., 2020; Zuo et al., 2020). Notably, it was found that SARS-CoV-2 infection could directly induce the release of NETs by healthy neutrophils (Veras et al., 2020). In line with these findings, we observed that neutrophils from acute patients released higher levels of DNA upon plasma stimulation. However, upon bacterial challenge the induction of NETs against SA was significantly reduced in acute COVID-19. This phenomenon might be due to neutrophil exhaustion and a subsequent inability to properly respond to bacterial challenges.

These findings were confirmed by the fact that neutrophils isolated from acute patients showed lower expression of key receptors CXCR1, CXCR2 as well as CXCR3, important for sensing IL-8 as well as G-CSF (Cummings et al., 1999; Swamydas et al., 2016). On the other hand, SDF-1α receptor CXCR4, involved in neutrophil trafficking from the bone marrow, was significantly higher, whereas the maturation marker CD15 was significantly lower in acute COVID-19 cells. Emergence of CXCR4+ cells has been linked as a neutrophil precursor marker (Evrard et al., 2018) and similarly it has been suggested that these immature neutrophils are being released into the blood during severe COVID-19 (Silvin et al., 2020). Also, presence of abnormal neutrophils in patients with severe COVID-19 has been observed (Wilk et al., 2020). A recent single-cell transcriptomic study proposed that premature neutrophils in severe COVID-19 might be programmed towards an anti-inflammatory start or even exert suppressive functions (Schulte-Schrepping et al., 2020). Our data here in addition underline the severity of the impairment of neutrophils’ ability to functionally respond to bacterial infection.

Acute patients showed a lower proportion of classical monocytes, crucial for anti-bacterial response, compared to rec-phase patient and healthy donors. Additionally, monocytes were characterized by lower numbers of non-classical monocytes that are important for maintaining vascular homeostasis (Giamarellos-Bourboulis et al., 2020a; Hadjadj et al., 2020; Schulte-Schrepping et al., 2020; Silvin et al., 2020; Thevarajan et al., 2020; Wilk et al., 2020). Similar to other studies, HLA-DR expression on classical monocytes was also significantly reduced (Giamarellos-Bourboulis et al., 2020a; Schulte-Schrepping et al., 2020; Silvin et al., 2020), which can be mediated by the IL-6 overproduction during severe COVID-19 (Giamarellos-Bourboulis et al., 2020a). Emergence of HLA-DR_low_ monocytes during severe COVID-19 can be linked to a phenotype similar to myeloid derived suppressor cells or dysfunctional monocytes. HLA-DR_low_, CD163_high_ monocytes are usually associated with anti-inflammatory tissue-homeostatic functions and are linked to an immunosuppressive phenotype in sepsis (Fischer-Riepe et al.; MacParland et al., 2018; Veglia et al., 2018; Venet et al., 2020). Thus, a defective or suppressed monocyte compartment can further add to its inability to respond to bacterial infection.

In conclusion, we demonstrated that during acute COVID-19, patients presented with alterations in neutrophil and monocyte effector cytokines, which severely affected their ability to respond to bacterial challenges. These data corroborated the clinical disease course with increased bacterial superinfections observed in critically-ill COVID-19 patients (Buehler et al., 2020). Our study further emphasizes the importance of tailoring treatments, aiming to restore the antibacterial effector functions of neutrophils and monocytes, thereby decreasing the risk of high lethality in COVID-19 due to secondary bacterial infections.

## Methods

### Human subject details

Patients recruited under the MicrobiotaCOVID prospective cohort study conducted at the Institute of Intensive Care Medicine of the University Hospital Zurich (Zurich, Switzerland) registered at clinicaltrials.gov (ClinicalTrials.gov Identifier: NCT04410263). The study was approved by the local ethics committee of the Canton of Zurich, Switzerland (Kantonale Ethikkommission Zurich BASEC ID 2020 - 00646). Patients were considered to be in the acute phase on the first 5 days after initial ICU admission while the recovery phase was defined as patients were discharged from the ICU or were negative for COVID-19 and under a defined clinical score, in a non-critical state. A list of all patient demographics and clinical scores is available in table S1 and S2.

### Bacterial strains

*Staphylococcus aureus* (SA) strains JE2 (MRSA-USA300, NARSA) and Cowan I (MSSA, ATCC 12598) were grown in Tryptic Soy Broth (TSB) at 37°C and 220rpm for 16 hours. Stationary phase cultures were diluted in fresh TSB and bacteria were grown to exponential phase for the infection. For *Streptococcus pneumoniae* (SP), the 603 strain (serotype 6B) (Malley et al., 2001) was passaged twice on blood agar plates (Columbia blood agar, Biomereux) and incubated at 37°C with 5% CO_2_ for 14h. A liquid culture was prepared in Todd Hewitt Yeast broth (THY) with a starting OD_600nm_ of 0.1 and grown at 37°C in a water bath until OD_600nm_ of 0.35 for the infection.

### Blood collection and plasma preparation

COVID-19 patients and healthy donors’ blood was sampled in EDTA tubes as per the protocol under the Microbiota-COVID (ClinicalTrials.gov Identifier: NCT04410263), (BASEC ID 2020 - 00646) and centrifuged at 3000 rpm for 10min for plasma collection. The collected plasma was centrifuged a second time at same conditions to remove any additional debris and supernatants were collected and aliquoted. Fresh plasma was immediately used to prepare a 10% plasma solution in RPMI 1640 (Gibco™) and used for *in vitro* experiments; the remaining plasma was utilized for cell stimulation as well as cytokine quantification and remaining aliquots stored at −80°C until further use.

### Cytokine measurement by Luminex

Cytokine levels in patients and healthy donors’ plasma, as well as cell culture supernatants from *ex vivo* experiments were assessed using the Luminex™ MAGPIX™ instrument (ThermoFisher). Samples were thawed on ice and prepared according to the manufacturer’s instructions using a custom-made 33-plex human cytokine panel (Procartaplex ThermoFisher). In brief, Luminex™ magnetic beads were added to the 96-well plate placed on a magnetic holder and incubated for 2min. The plate was washed twice with assay buffer for 30sec. In parallel, provided standards and plasma samples were diluted in assay buffer (cell culture media was used for cell culture supernatants) and added to the plate. The plate was incubated for 2h at RT at 550rpm in a plate orbital shaker. Next, the plate was washed twice with assay buffer and incubated for 30min at 550rpm with detection antibodies. After two washing steps, the plate was incubated with Streptavidin-PE solution for 30min at 550rpm. Finally, the plate was washed, reading buffer was added and incubated for 10min at RT and 550rpm before running the plate. Data acquisition and analysis were performed using the Xponent software (v. 4.3). Data were validated using the Procarta plex analyst software (ThermoFisher).

### Principal Component and Integrated Correlation analysis

PCA plots of the cytokine analysis from patient and healthy donor plasma as well as receptor analysis from the ex vivo experiments were created using the ‘PCA’ and the ‘fviz_pca_biplot’ functions available in ‘FactoMineR’ package in R. Correlation mapping was performed using the ‘corrplot’ package in R. The color of the circles indicated positive (blue) and negative (red) correlations, color intensity represented correlation strength as measured by the Pearson’s correlation coefficient. The correlation matrix was reordered manually to better visualize the variables of interest.

### Peripheral blood mononuclear cells (PBMCs) isolation

Patients and healthy donor PBMCs were isolated from the cellular fraction of the blood after 1:2 dilution with DPBS using the Lymphoprep (Axis Shield) density gradient method. In brief, the diluted blood was overlaid on Lymphoprep and centrifuged for 25min at 2000rpm with lowest acceleration and break settings. Following the gradient separation, the PBMCs layer was transferred into a new 50ml conical tube and diluted with FACS buffer (2mM EDTA and 1% FBS). Cells were washed twice with FACS buffer. Next, cells were resuspended in red blood cells (RBC) lysis buffer (ThermoFisher), mixed gently and incubated for 10min at 37°C and 5% CO_2_. The lysis reaction was stopped by adding FACS buffer and the suspension was centrifuged. Cells were washed once and resuspended in FACS buffer for counting using the Attune NxT flow cytometer (ThermoFisher).

### Monocytes enrichment from PBMCs

Patient and healthy PBMCs were used for monocyte enrichment using the EasySep™ Human Monocyte Enrichment Kit without CD16 Depletion (StemCell™) following the manufacturer’s instructions. In brief, PBMCs (<100 million) were transferred to a 5ml polystyrene tube, the human monocyte enrichment cocktail was added and the sample was gently mixed and incubated for 10min on ice. Following incubation, magnetic beads were added to the mixture and samples were mixed and incubated for 10min on ice. Finally, the mixture was placed in a magnetic holder (StemCell™) for 3min and the cells were decanted into a new tube. Monocytes were then washed, resuspended in RPMI 1640 and counted using the Attune NxT flow-cytometer (ThermoFisher).

### Neutrophils isolation

Neutrophils were isolated from the cellular fraction of the blood, after dilution with DPBS (Gibco™), with the EasySep™ Human Neutrophil Isolation Kit (StemCell™) according to the manufacturer’s instruction. In brief, Neutrophil enrichment cocktail was added to the diluted blood and incubated for 15min at RT. Next, magnetic beads were added for another 15min, after which the tubes were placed into a magnetic holder (StemCell™). PMNs were collected after 15min of cell separation. They were centrifuged at 1500rpm for 6min (low acceleration and brakes) and subsequent red blood cells (RBCs) lysis was performed with resuspension in H2O followed by addition of DPBS. After a further centrifugation step, neutrophils were resuspended in RPMI 1640 and counted using the Attune NxT flow-cytometer (ThermoFisher).

### Plasma stimulation

Isolated neutrophils or monocytes, from both COVID-19 patients and healthy donors, were seeded in conical 96-well V-bottom plates (for Flow cytometry assays, around 2×10^5^ cells / well) or in 24-well F-bottom plates (for phagocytosis and intracellular survival assays, approximately 2.5×10^5^ to 3×10^5^ cells/well) and stimulated with 10% autologous or heterologous (either COVID-19 or healthy donor plasma) for 2.5h at 37°C + 5% CO_2_.

### Bacterial challenge

For phagocytosis and intracellular killing assays, bacteria were opsonized for 20min in RPMI 1640 supplemented with 2.5% of either patient or healthy plasma at a determined multiplicity of infection (MOI) for neutrophils (50 for SP and 10 for JE2) and monocytes (50 for SP and Cowan I) infections respectively.

To analyze intracellular survival of the bacteria in neutrophils, neutrophils were seeded into 24-well plates (TPP) and infected with exponentially grown SA at a MOI of 10 or with exponentially grown SP at a MOI of 50. After 40min, 1mg/ml flucloxacillin and 25μg/ml lysostaphin were added to kill all extracellular SA or penicillin (10μg /ml) / streptomycin (10μg/ml) to kill extracellular SP. Infected cells were harvested 30min and 4h after addition of antibiotics, washed twice with PBS, lysed with ddH_2_O, serially diluted and drop plated. Bacterial survival was analyzed and calculated relatively to the invasion (30min time point).

To analyze intracellular survival of the bacteria within monocytes were seeded into 24-well plates (TPP) and infected with exponentially grown SA or SP at a MOI of 50. After 40min, 1mg/ml flucloxacillin and 25μg/ml lysostaphin were added to kill all extracellular SA or penicillin (10u/ml) / streptomycin (10μg/ml) to kill extracellular SP. Infected cells were harvested 30min and 90min after addition of antibiotics, washed twice with PBS, lysed with 0.02% of Triton X-100 in ddH_2_O, serially diluted and drop plated. Bacterial survival was analyses and calculated relatively to the invasion (30min time point). End point bacterial free supernatant from both neutrophils and monocytes bacterial infection experiments were utilized for cytokine measurement.

### Flow cytometry

Staining of reactive oxygen species (ROS, for neutrophils and monocytes) and nitric oxide (NO, formonocytes) was performed after 1 hour of bacterial challenge or plasma stimulation only by incubation with 5μM CellROX™ green reagent and 5μM DAF-FM™ diacetate (ThermoFisher) respectively, for 30min. After the incubation period, cells were washed with DPBS. Cells were stained with either LIVE/DEAD™ fixable Near-IR or Aqua stain (ThermoFisher) in DPBS for 25min at 4°C. Next, cells were washed with FACS buffer and stained for surface antigens for 30min at 4°C. Antibodies included anti-CD15 eFluor450 (clone: HI98), anti-CD181 FITC (8F1-1-4), anti-CD182 PerCP-eFluor710 (5E8-C7-F10), anti-CD183 PE-eFluor610 (CEW33D), anti-CD66b APC (G10F5), anti-HLA-DR eFluor450 (LN3), anti-CD45 eFluor506 (HI30), anti-CD14 SB600 (61D3), anti-CD64 FITC (10.1), anti-CD163 PerCP-eFluor710 (GHI/61), anti-CD16 PE (CB16), anti-CD86 PE-Cyanine 5.5 (IT2.2), anti-CD206 PE-Cyanine 7 (19.2), anti-CD169 APC (7-239), anti-CD11b AF® 700 (VIM12), anti-CD3 APC-eFluor780 (UCHT1), anti-CD19 APC-eFluor780 (HIB19), anti-CD56 APC-eFluor780 (CMSSB), anti-CD119 FITC (BB1E2) and anti-CX_3_CR1 APC (2A9-1) from ThermoFisher, anti-CD195 BV510 (J418F1), anti-CD184 BV605 (12G5), anti-CD191 PE-Cyanine7 (5F10B29), anti-CD88 AF®700 (S5/1), anti-CD192 PerCP-Cyanine5.5 (K036C2), anti-CD80 PE-Dazzle™594 (2D10) and anti-CD120b PE-Cyanine7 (3G7A02) from Biolegend. For intracellular MPO staining, neutrophils were washed, fixed and permeabilized with the Cytofix/Cytoperm™ Fixation/ Permeabilization Solution Kit (BD) for 15min at 4°C and stained subsequently for another 30min with anti-MPO eFluor450 (455-BE6). To assess neutrophil extracellular traps (NETs), cells were stained first with LIVE/DEAD™ fixable Aqua, followed by staining for surface antigens as described above, after which they were washed with DPBS and subsequently stained with SYTOX™ Green in DPBS for 30min. To stain for extracellular MPO-DNA complexes, neutrophils were stained exactly as described for NETs with the addition of the MPO staining during the surface antigen step. Cells were analyzed on an Attune NxT (ThermoFisher). All antibodies and concentrations used are listed in Table x. Flow cytometry data were analyzed with FlowJo (v10.2). Neutrophils and monocytes were gated based on their forward- and side-scatter properties, single cells and ultimately live cells. Neutrophils were characterized as CD66b+CD16+, whereas monocytes were divided into subgroup based on CD14+CD16- (classical), CD14+CD16+ (intermediate) and CD14dimCD16+ (non-classical) for further analysis.

### Microscopy and NETs quantification

Neutrophils were stimulated as described above and placed within wells of a μ-slide (iBidi) and centrifuged at 200 g for 2 min, after which they were challenged with *S. aureus* for 1.5h. NETs were stained by directly adding SYTOX™ Green and 2μM Hoechst 33342 (ThermoFisher) for 30min at room temperature to the wells. The confocal laser scanning microscopy images were obtained with a Leica TCS SP8 inverted microscope using a 63×/1.4 oil immersion objective. The whole wells were inspected for NETs formation and two to three representative spots per condition were imaged. The obtained images were processed using Imaris 9.2.0 software (Bitplane) to obtain tifs for further analysis. Other standard light microscopy images of fixed cells were obtained on a fully automated Olympus IX83 with a 40X objective (UPLFLN40XPH-2) illuminated with a PE-4000 LED system through a quadband filter set (U-IFCBL50). 16 positions per sample were assigned before the sample was prepared to avoid potential experimenter bias. Automated NET quantification was performed as described in SI Fig.4: after filtering nuclei on DAPI signal (threshold set manually for each 8-samples experiment), extracellular DNA was quantified on Sytox Green signal. Images containing large cell aggregates that could not be resolved were discarded. Nuclei were counted after watershed segmentation on the DAPI mask. Images were processed using ImageJ software (Rasband, W.S., ImageJ, U. S. National Institutes of Health, Bethesda, Maryland, USA, https://imagej.nih.gov/ij/, 1997-2018) and Matlab R2020a (MathWorks).

### Statistical analysis

The number of donors is annotated in the corresponding figure legend. Differences between two groups were evaluated using either Mann-Whitney test or Wilcoxon signed-rank test. Kruskal-Wallis test with Dunn’s multiple comparisons test was used to evaluate differences among the three groups in all the analyses (GraphPad). Pearson test was used for correlations of normally distributed binary data. Significance level with p<0.05 are depicted in individual graphs.

For the statistical analyses involving several cytokines, measured cytokine values were normalized based on the standard z-score formula. This allowed to compare cytokines to each other and to obtain a sum of z-scores per patient.

## Acknowledgments

Confocal laser scanning microscopy was performed with the support of the Center for Microscopy and Image Analysis, University of Zurich. We thank Andrea Tarnutzer and Federica Andreoni for their technical help and inputs on the manuscript.

## Funding

This work was funded by the SNSF project grant 31003A_176252 (to A.S.Z), the SNF Biobanking grant 31BK30_185401 (to A.S.Z), the Uniscientia Foundation Grant (to A.S.Z), by the Swedish Society for Medical Research (SSMF) foundation grant P17-0179 (to S.M.S), the Promedica Foundation 1449/M (to S.D.B) and unrestricted funds (to RAS).

The MicrobiotaCOVID prospective cohort study was approved by the local ethics committee of the Canton of Zurich, Switzerland (Kantonale Ethikkommission Zurich BASEC ID 2020 - 00646)

## Author Contributions

**Conceptualization:** SMS, SDB and ASZ

**Investigation:** SMS, AGM, TAS, SDB and ASZ

**Experimental design:** SMS, AGM, TAS, MH and ASZ

**Methodology:** SMS, AGM, TAS, MH, CCC, CA, CV, NM, DAH, PBK and SDB

**Data curation:** SMS, AGM, TAS, MH, CV, CA, JBP, MPH, PDWG and SDB

**Formal Analysis:** SMS, AGM, TAS, MH, CA, JBP, and CV

**Funding acquisition:** SDB, PBK, RAS, and ASZ

**Visualization:** SMS, AGM, TAS, JBP and CV

**Resources:** PBK, SDB, RAS and ASZ

**Writing – original draft:** SMS, AGM, TAS and MH

**Writing – review & editing:** SMS, AGM, TAS, MH, CCC, CA, JBP, CV, DAH, PBK, RAS, SDB and ASZ

## Declaration of Interests

The authors declare no competing interests

## Notes

### Competing Interest Statement

The authors have declared no competing interest.

