## Supplementary Table and Figures for "Neutrophil and monocyte dysfunctional effector response towards bacterial challenge in critically-ill COVID-19 patients"

**Original article:**

**Key words: COVID-19, Neutrophils, Monocytes, secondary bacterial infections, hypercytokinemia**

### Supplementary Tables and Figures

Table S1: Clinical characteristics of the patients

| Clinical Characteristics | Study Patients (n=27) |
|---|---|
| Age (mean ± SD) | 59.9 (±8.4) |
| Male (%) | 23 (85%) |
| Diabetes (%) | 13 (48%) |
| COPD/Asthma (%) | 3 (11%) |
| Charlson Comorbidity Index (mean ± SD) | 1.7 (1.9) |
| Malignancy (%) | 3 (11%) |
| Immunosuppression (%) | 4 (15%) |
| 28 Day Survival (%) | 24 (89%) |
| Alive at ICU Discharge (%) | 22 (81%) |
| Pulmonary Superinfection (%) | 15 (56%) |
| Bacteremia (%) | 10 (37%)** |
| Length of Mechanical Ventilation (mean ± SD) | 21.9 (±21.2) |
| Length of ICU-Stay (mean ± SD) | 28.4 (±26.7) |
| SOFA Score Day 1 (mean ± SD) | 7.5 (±3.2) |
| SAPS on Admission (mean ± SD) | 39.9 (±16.5) |

Table S2: Treatment received by the patients

| Patient-ID | Acute phase (n=27) | | | | Recovery phase (n=21) | | | |
|---|---|---|---|---|---|---|---|---|
| | Anti-infectives | Hydroxy-chloroquin | Steroids | Other Immuno-suppression | Anti-infectives | Hydroxy-chloroquin | Steroids | Other Immuno-suppression |
| 1 | Yes | No | No | No | - | - | - | - |
| 2 | Yes | Yes | Yes | Yes | Yes | No | Yes | Yes |
| 3 | Yes | No | No | No | No | No | No | No |
| 5 | Yes | Yes | Yes | No | Yes | No | No | No |
| 7 | Yes | No | No | Yes | - | - | - | - |
| 8 | Yes | No | Yes | Yes | Yes | No | Yes | Yes |
| 9 | Yes | No | No | No | Yes | No | No | No |
| 10 | Yes | No | No | No | Yes | No | Yes | No |
| 11 | Yes | No | No | No | Yes | No | No | No |
| 12 | Yes | No | Yes | No | Yes | No | Yes | No |
| 13 | Yes | No | No | No | No | No | No | No |
| 14 | Yes | No | Yes | No | - | - | - | - |
| 15 | Yes | No | Yes | No | No | No | No | No |
| 16 | Yes | No | Yes | No | Yes | No | Yes | No |
| 17 | Yes | No | Yes | No | - | - | - | - |
| 18 | Yes | No | Yes | No | Yes | No | Yes | No |
| 19 | Yes | No | Yes | No | No | No | Yes | No |
| 22 | Yes | No | Yes | No | - | - | - | - |
| 23 | Yes | No | Yes | No | - | - | - | - |
| 127 | Yes | Yes | No | No | Yes | No | Yes | No |
| 132 | Yes | Yes | No | No | Yes | No | No | No |
| 134 | Yes | No | Yes | No | No | No | No | No |
| 140 | Yes | Yes | Yes | Yes | No | No | No | Yes |
| 143 | Yes | No | No | No | Yes | No | Yes | No |
| 153 | Yes | No | No | No | No | No | No | No |
| 154 | Yes | Yes | No | No | Yes | No | Yes | No |
| 157 | Yes | Yes | No | No | Yes | No | No | No |
| Total | 27/27 (100%) | 7/27 (26%) | 14/27 (52%) | 4/27 (15%) | 14/21 (67%) | 0/21 (0%) | 10/21 (48%) | 3/21 (14%) |

<u>Definitions:</u>

Anti-infective: Use of either/or antibiotics, antivirals, antifungals in the last 48 hours before sampling

Hydroxychloroquin: Use of hydroxychloroquin in the last 7 days before sampling

Steroids: Any use of steroids equivalent to 20mg prednisone per day in the last 48 hours before sampling

Other Immunosuppression: Any use of other immunosuppression equivalent to 20mg prednisone per day in the last 48 hours before sampling

**Supplementary Figure 1**

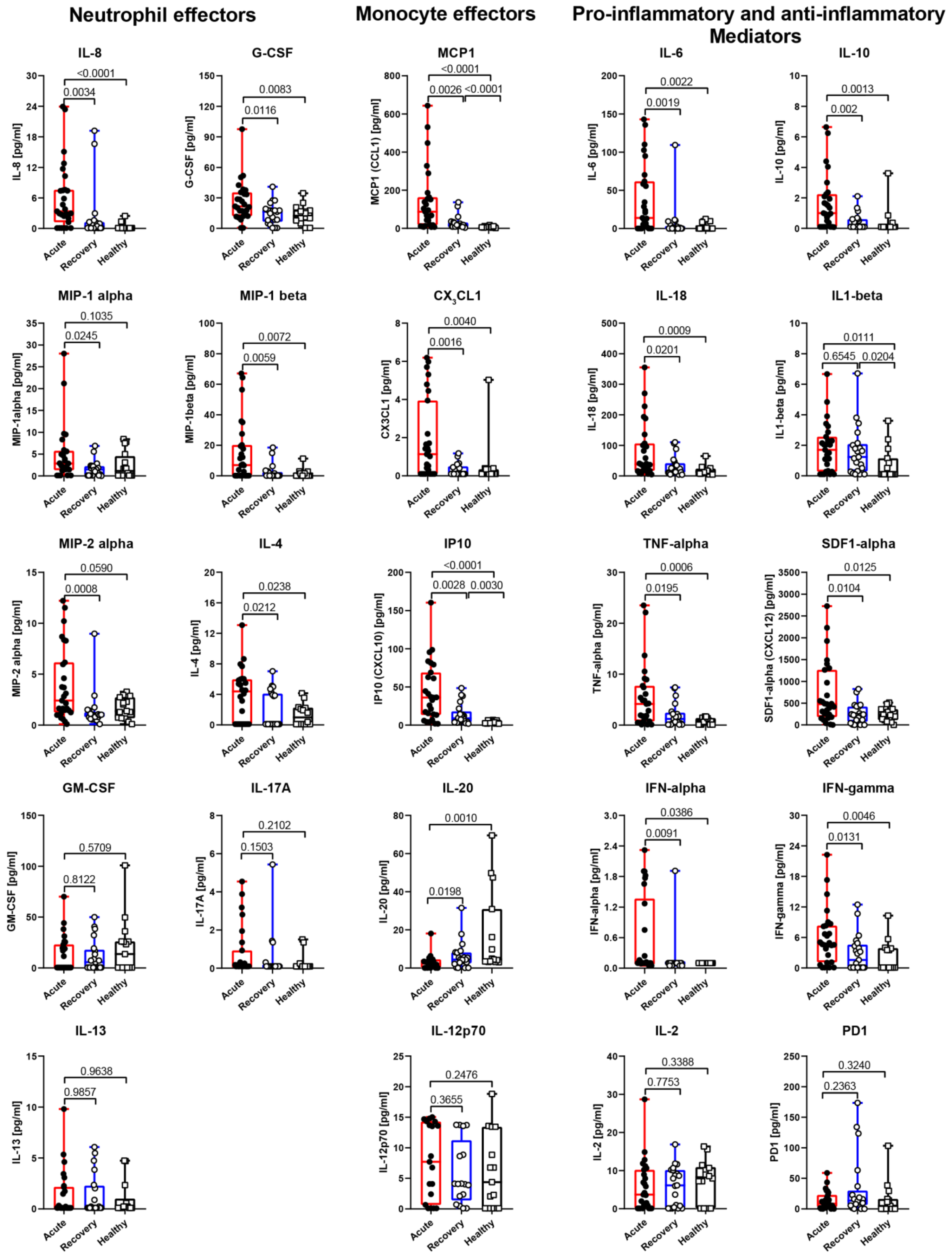

**Figure S1**. Cytokine levels in plasma from COVID-19 patients in acute (n=27) or recovery (n=21) phase as well as healthy donors (n=16). 24 different cytokines, grouped as neutrophil effectors, monocyte effectors and pro-inflammatory and anti-inflammatory mediators, were determined from plasma using a luminex multiplex assay.

### Supplementary figure 2

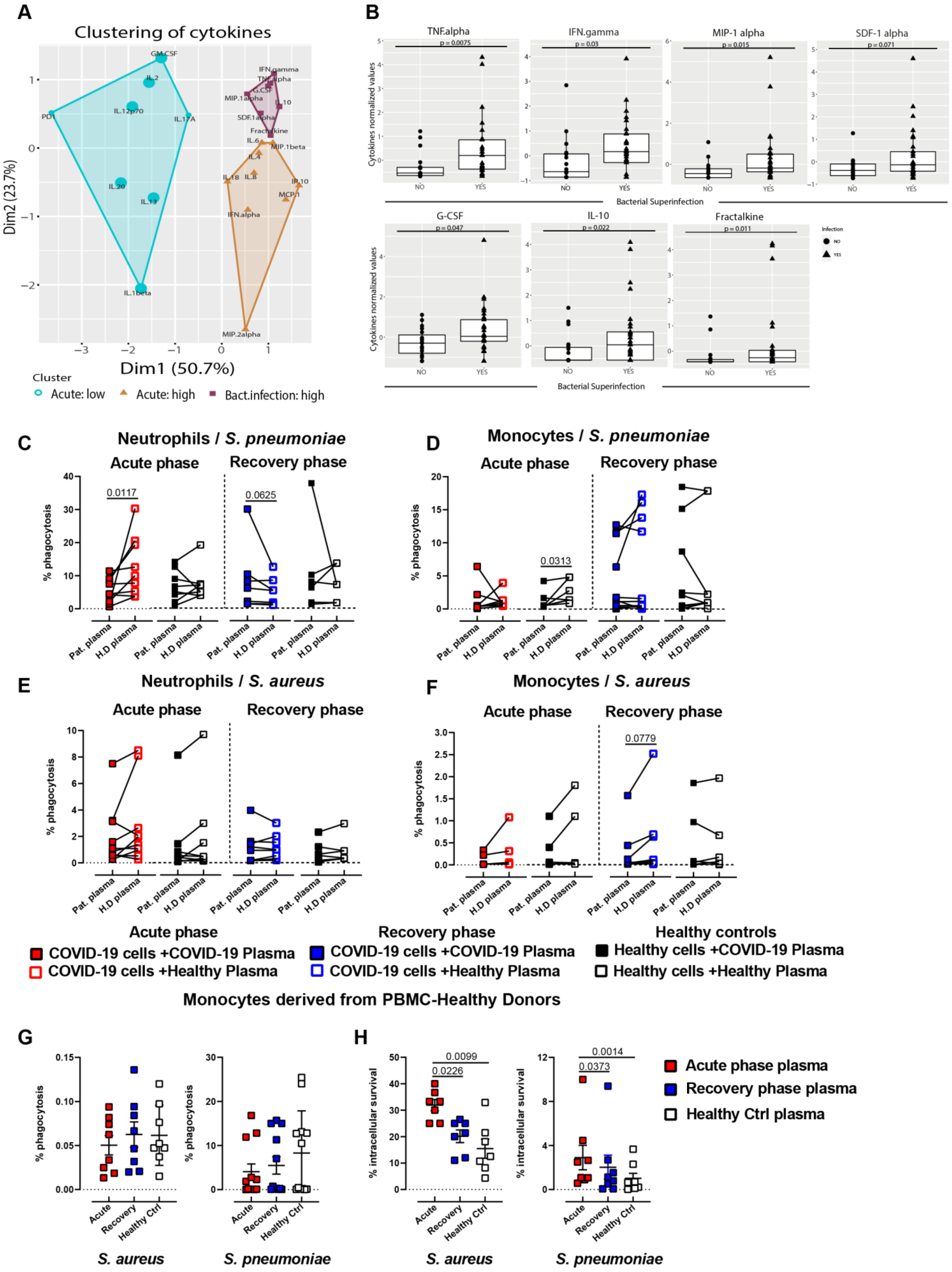

**Figure S2.**

(**A**) Principal component analysis of cytokine levels in COVID-19 patients with secondary bacterial infection versus COVID-19 patients without secondary bacterial infection. Cytokine cluster exhibiting elevated levels in patients with secondary bacterial infection (both acute and rec phase; purple), cluster with elevated levels in patients without superinfection (only acute phase; orange) and cluster with similar levels between acute and rec. phase in patients without superinfection (cyan). (**B**) Normalised cytokine values (sum of Z-scores) in the plasma of acute and recovery patients with or without bacterial superinfection**.** Phagocytosis capacity of COVID-19 patient neutrophils (left) and monocytes (right) pre-exposed to plasma from patients (solid symbols) vs healthy plasma (open symbols) upon infection with SP (**C - D**) or SA (**E - F**). COVID-19 monocyte phenotype induction using healthy PBMCs upon acute, rec-phase and healthy plasma exposure and its ability to phagocyte (**G**) and clear intracellular SA or SP (**H**).

Supplementary Figure 3

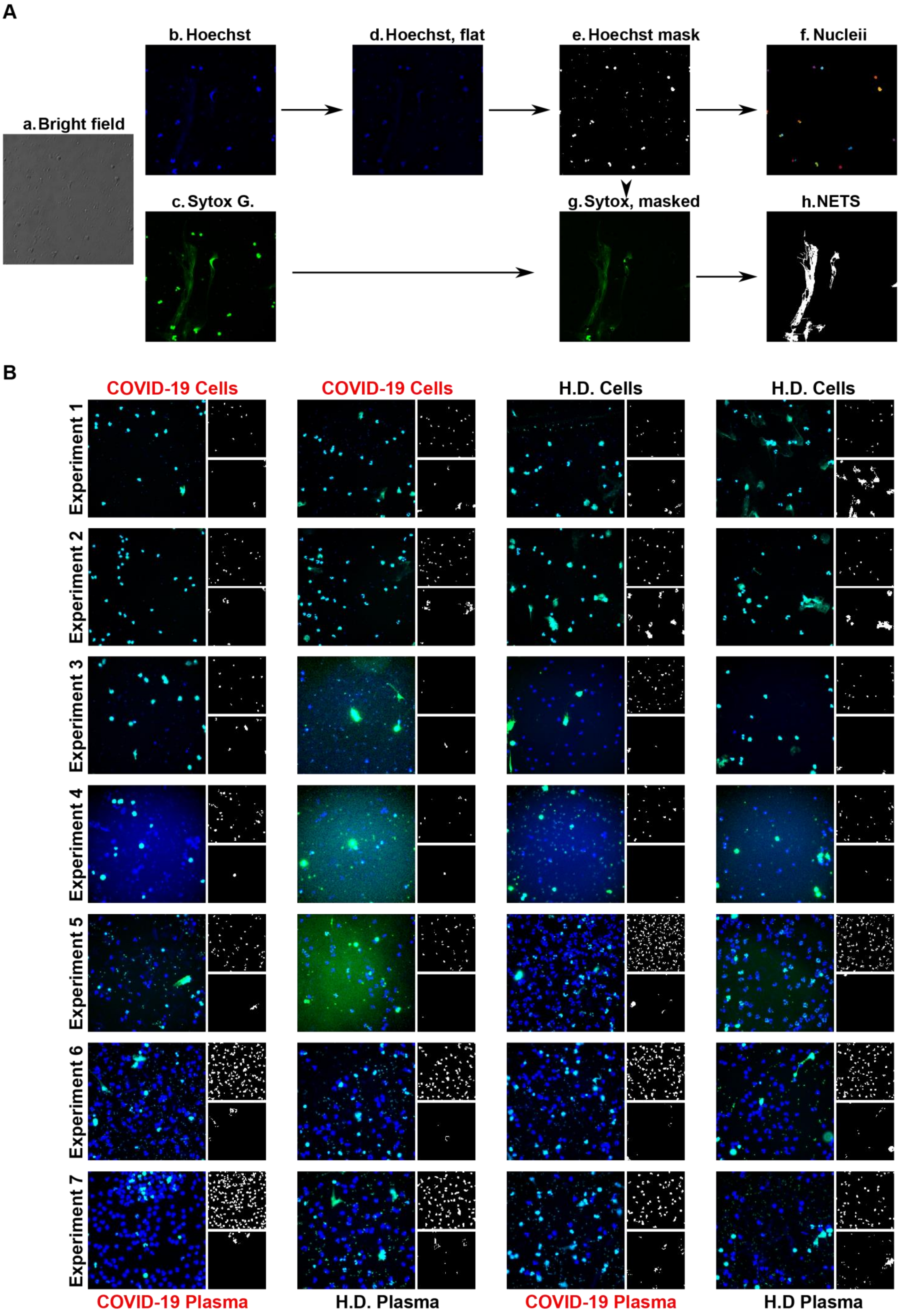

**Figure S3**. **Image analysis process used to quantify NETs from microscopy images** (**A**, sub categories **a, b, c**) using Hoechst and Sytox green images. (**d)** First flatfield correction is applied to Hoechst images in order to get rid of vignetting. (**e**) A mask is then obtained through thresholding and dilating the binary image obtained. **(f).** Nuclei are then quantified after filtration of the small objects resulting from autofluorescence of plasma particles and *S. aureus* cells and watersheding (**g).** The NETs mask is obtained after applying the Hoechst mask to the Sytox Green images, and then thresholding and filtering out small objects (**h**). **Representative images for quantified NETs using microscopy images (B)** Each COVID 19 patient / healthy donor (H.D.) control pair is shown on a line with the 4 possible plasma / cells permutations. Each montage of 3 pictures is issued from the most representative image in terms of NET area per Nuclei values, out of the 16 images obtained per sample, at randomized positions. The montage consists in the Hoechst and Sytox green overlay (**left**) and the resulting nuclei mask (**top right**) and the obtained mask (**bottom right**).

### Supplementary Figure 4

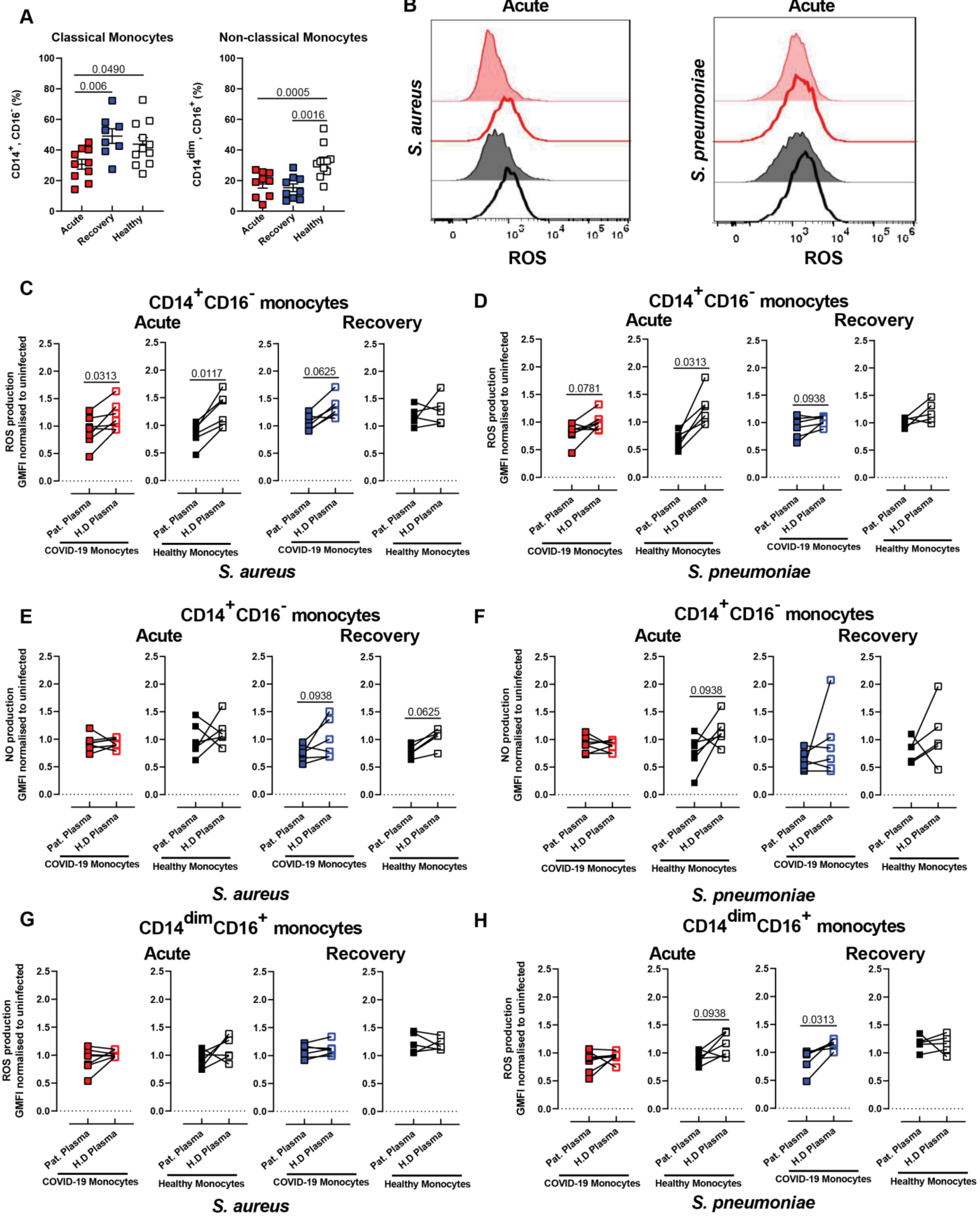

**Figure S4**. Characterization of COVID-19 monocyte sub-type distribution based on their cell surface expression of CD14 and CD16 (**A**) and Representative histogram of the ability of monocytes to produce ROS (**B**) and normalized GMFI values (**C – D and G - H**), as well as nitric oxide (NO) production (**E – D**) upon induction with COVID-19 patient plasma and following challenge with SA or SP.

### Supplementary Figure 5

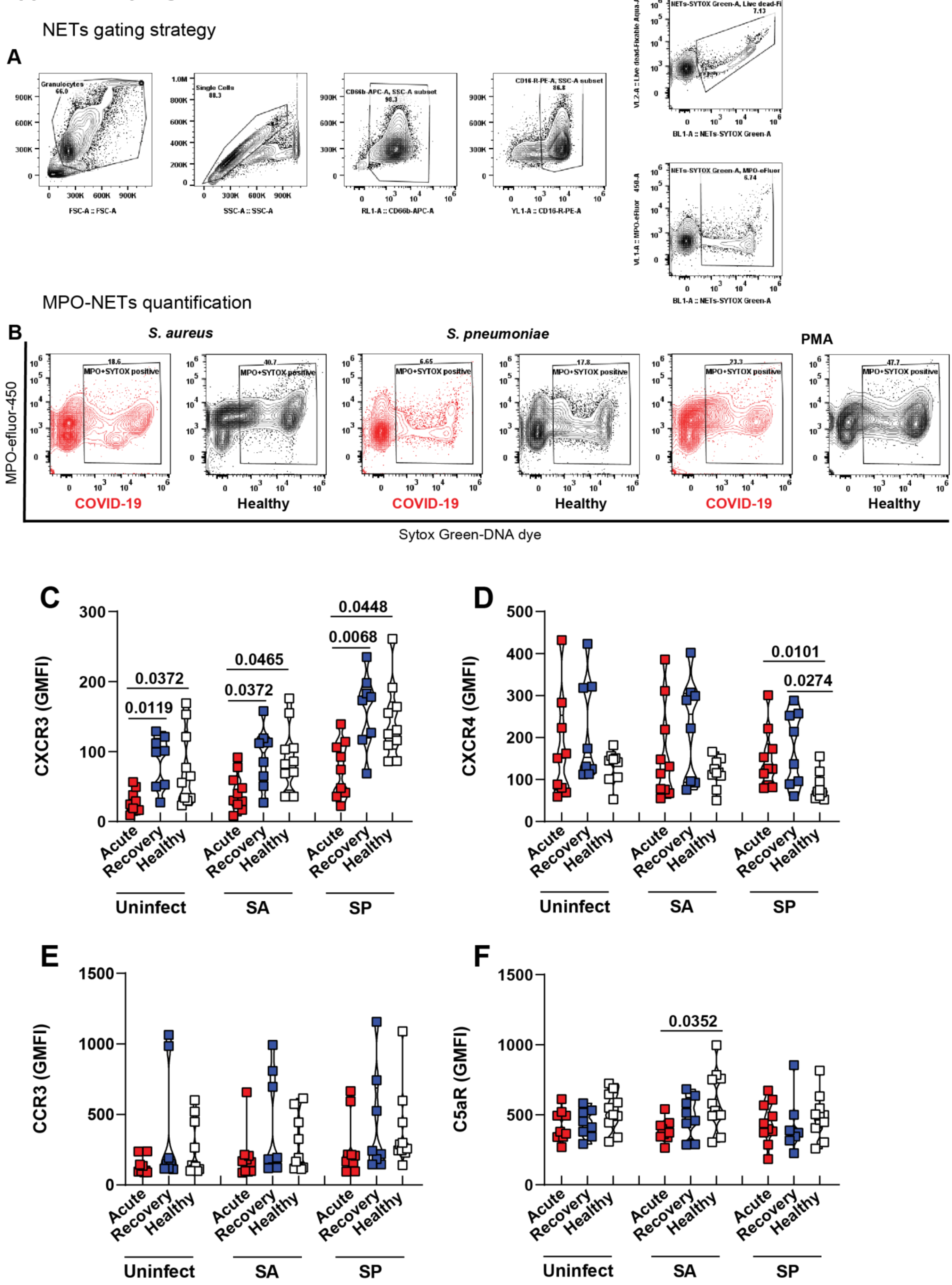

**Figure S5.** (A--B) Gating strategy for the quantification of NETs production in neutrophils by Flow-cytometry using the cell markers: $CD66b^+$, $CD15^+$, Live-dead fixable $Aqua^-$ and SYTOX $green^+$ - $MPO^+$. Characterization and quantification of surface receptors CXCR3 (**A**), CXCR4 (**B**), CCR3 (**C**) and C5aR (**D**) in COVID-19 neutrophils upon stimulation with either COVID-19 patient plasma (red-acute or blue-recovery) or healthy plasma (white) and further challenge with SA or SP.

### Supplementary Figure 6

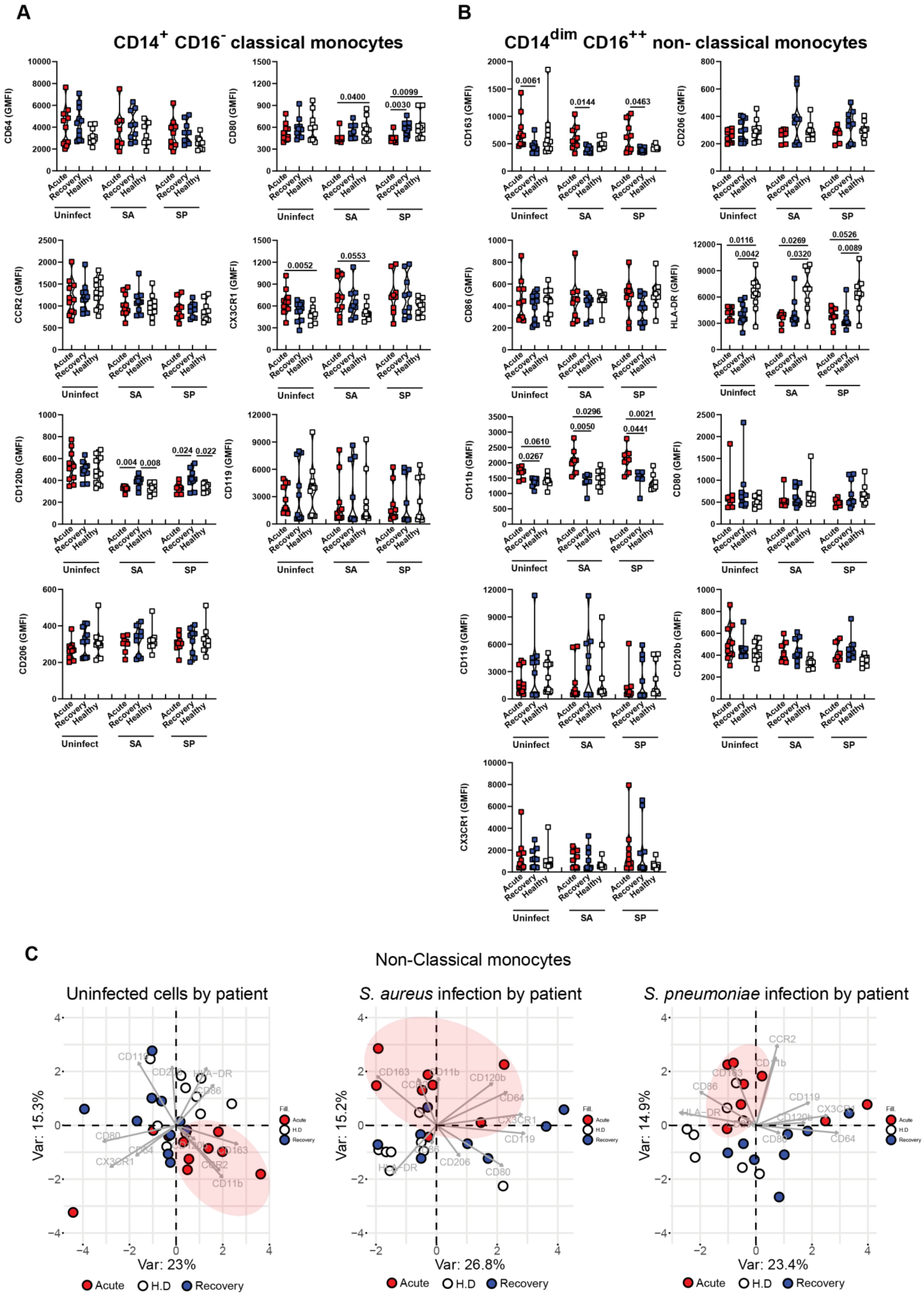

**Figure S6.** Determination of the expression of key surface markers in COVID-19 acute (red), recovery (blue) as well as in healthy donors (white) $CD14^{+}$ $CD16^{-}$ classical monocytes (**A**) and $CD14^{dim}$ $CD16^{++}$ non-classical (**B**) monocytes using flow-cytometry. PCA plot of cell surface phenotype of COVID-19 patients acute (red), recovery (blue) as well as in healthy donors (white) for the non-classical monocytes (**C**) from COVID-19 basal level without bacterial challenge, upon SA infection, or SP infection.
